# Larval mannitol diets increase mortality, prolong development, and decrease adult body sizes in fruit flies (*Drosophila melanogaster*)

**DOI:** 10.1101/674192

**Authors:** Meghan Barrett, Katherine Fiocca, Edward A. Waddell, Cheyenne McNair, Sean O’Donnell, Daniel R. Marenda

**Author notes:** These authors contributed equally to this work (Meghan Barrett, Katherine Fiocca). These authors also contributed equally to this work (Sean O’Donnell, Daniel R. Marenda). Correspondence author; (DRM).

## Abstract

Ingestion of the polyol mannitol caused sex-biased mortality in adult *Drosophila melanogaster*, but larval mortality was not sex-biased. High-sugar diets prolong development and generate smaller adult body sizes in *D. melanogaster*. We hypothesized that mannitol ingestion would generate similar developmental phenotypes as other high-carbohydrate diets. We predicted concentration-dependent effects on development similar to high-sugar diets when *D. melanogaster* larvae are fed mannitol, as well as a concentration-dependent amelioration of developmental effects if introduction to mannitol media is delayed past the third instar. Both male and female larvae had prolonged development and smaller adult body sizes when fed increasing concentrations of mannitol. Mannitol-induced increases in mortality were concentration dependent in 0 M to 0.8 M treatments beginning as early as 48 hours post-hatching. Larval survival, and pupation and eclosion times, were normal in 0.4 M mannitol treatments when larvae were first introduced to mannitol 72 hours post-hatching (the beginning of the third-instar); the adverse mannitol effects occurred in 0.8 M mannitol treatments, but at a lower magnitude. Female *D. melanogaster* adults prefer laying eggs on diets with high sugar concentrations, despite the negative effects on offspring performance. However, when given a choice, female *D. melanogaster* avoided laying eggs on mannitol-containing media that was otherwise identical to the control media, suggesting females perceived and avoided mannitol. In conclusion, the developmental effects of a larval mannitol diet closely resemble those of high-sugar diets, but adult female oviposition responses to mannitol in laying substrates are distinct from responses to other carbohydrates.

## Introduction

Developmental duration and body size are controlled by three related variables in holometabolous insects: growth rate, critical weight (the point at which the developmental period is no longer affected by resource levels), and the interval to the cessation of growth [1,2]. Because they are controlled by the same three parameters, a direct, positive relationship is expected and typically observed between developmental duration and body size [3–8].

However, some environmental variables can differently affect growth rate, critical weight, and interval to the cessation of growth, causing neutral or even negative relationships to occur between body size and development time [2,4]. High-carbohydrate diets, specifically sucrose and glucose, affect insect growth and development. High-sugar diets disrupt the insulin/TOR signaling pathway through increased circulating trehalose levels [9,10]. High-sugar fed *D. melanogaster* adults are a model system for studying metabolic phenotypes associated with insulin resistance and diabetes [9–15]. At the larval stage, high-sugar diets lead to delays in adult eclosion (due to delayed onset of pupation, but not prolonged pupation periods), reduced survival, and smaller pupal case volumes and lower adult dry mass [10,16–19].

Mannitol, a non-sugar polyol carbohydrate, prolonged development when fed to *D. melanogaster* larvae [20], and larvae fed mannitol were smaller than control larvae of the same age (Fiocca and Barrett, personal observation). Mannitol is a sugar alcohol and isomer of sorbitol. It is produced naturally as a product of fermentation and is found commonly in plants, bacteria, and fungi [21–23]. Mannitol is used as a low-calorie sweetener, sweetening foods without increasing blood glucose levels or insulin in humans [24,25]. However, ingestion and breakdown of mannitol by *Tribolium castaneum* beetles increased hemolymph trehalose levels, indicating mannitol may be a nutritive source of dietary carbohydrates in some insect taxa [26,27]. We hypothesized that mannitol ingestion during *D. melanogaster* development would generate phenotypes similar to those produced by high sugar diets [10,17,18]. The ability of polyols to disrupt development has not been studied, and identifying additional compounds that affect insect development can further our understanding of the pathways that connect growth rate, developmental timing, and body size in insects.

The timing of high-carbohydrate diet introduction to larvae is important in determining its effects, particularly pre- and post-critical weight [10,28]. Third instar larvae fed a high-sugar diet showed lower transcriptional changes in the expression of genes associated with glucose transport and metabolism, lipid synthesis and storage, trehalose synthesis and stability, and oxidative stress when compared to first instar larvae continuously fed sucrose [10]. We hypothesized that delaying mannitol introduction to larvae until the third instar would reduce the severity of mannitol’s developmental effects in larvae fed high molarity mannitol media.

Female *D. melanogaster* choose high-carbohydrate (sucrose) oviposition sites, even when these sites detrimentally affect the fitness of their offspring [16]. However not all carbohydrates induce this same response; notably, high-carbohydrate erythritol substrates did not affect oviposition choice compared to lower-carbohydrate substrates [29]. The impact of mannitol, found in both fresh and rotting fruits due to microbial fermentation, on oviposition choice has not been explored [23,30].

In this study, we quantified the effect of mannitol feeding as a larva on adult body size, measured by thorax length. We assessed the effects of increasing concentrations of dietary mannitol on *D. melanogaster* larval survival, and pupation and eclosion times. We analyzed if developmental delays were due to a delay in the onset of pupation, and/or prolonged time in the pupal stage. We also evaluated if delaying mannitol introduction to larvae by 72 hours, or approximately the early third instar [31], could reduce or eliminate the developmental effects of decreased survival and prolonged developmental duration. We assessed if adult females differed in the preference for control vs. mannitol media for oviposition sites. Mannitol ingestion during the larval stage is a rare example of environmental substrate variation that can decouple the typical positive relationship between development duration and body size; the effects were concentration-dependent and developmental stage-dependent. We discuss the similarities between larval mannitol diets and high-sugar diets, and hypothesize that the insulin signaling pathway is a possible mechanism for mannitol’s developmental effects.

## Methods and materials

### Culturing *Drosophila*

Wild-type (Canton S) *D. melanogaster* (Bloomington *Drosophila* Stock Center) were raised to adulthood on standard *Drosophila* media for laboratory culturing and reared in an insect growth chamber at 27.5 °C, 50 % relative humidity, with a 12-h:12-h photoperiod [32]. These conditions were used to rear adults and for all larval experiments. Standard media was prepared in 100 ml batches as follows: 9.4 g cornmeal, 3.77 g yeast, 0.71 g agar, 0.746 ml Propionic acid, 1.884 ml Tegosept (10 % w/v methyl p-hydroxybenzoate in 95 % ethanol), and 9.42 ml molasses (Genesee Scientific). The appropriate amount of mannitol (HiMedia; GRM024-500G, Lot 000249743) was added, and beakers were filled with distilled water to a final volume of 100 ml. After heating the mixed ingredients to set the agar, media was poured into vials and cooled until consistency was firm and uniform. An excess of media was provided, with 10 ml in each vial.

### Testing effect of larval mannitol feeding on adult body size

Groups of 15 male and 15 female wild-type flies raised on standard media were placed in vials containing 0 M, 0.4 M, or 0.8M mannitol adult media (standard media recipe with no molasses) and allowed to lay for 24 hours (at which time they were removed). Nine vials were used per concentration, with a total of 405 flies of each sex. Vials were checked for newly emerged adults every twelve hours from Day 10 to Day 15, and every twenty-four hours from Day 15 to Day 24 (the last day that a larva pupated in the larval plate trials). Adult flies were removed from the vials and sexed; two males and two females were randomly selected every 24 hours from each vial with adults. Selected adults were sacrificed and photographed for body size measurements (0M: n= 52 females, n= 56 males; 0.4M: n= 66 females, n= 61 males; 0.8M: n= 50 females, n= 49 males). Photographs of the thorax were taken from a dorsal view at 4 X magnification using a digital camera mounted (0.7 X) on a dissecting scope. Measurements of thorax length were taken from the tip of the scutellum to the most anterior part of the mesothorax [33,34] in ImageJ using the ruler tool [35], and photographs of a stage micrometer were used to convert pixels to mm.

### Testing effects of dietary mannitol on larval mortality and developmental delay

Translucent media was produced by omitting the cornmeal from the standard media recipe and lowering the amount of agar to 0.52 g/100 ml [29]. Food was poured to a depth of 3 mm in 50 mm diameter petri dishes, allowing for the observation of the larvae in the food. Groups of over 100 mixed male and female wild-type flies raised on standard media were placed in each of 10 egg laying chambers. At the end of four hours, eggs were collected and five eggs were plated per petri dish, with mannitol concentrations from 0 to 0.8 M, at 0.2 M increments. Six petri dishes were used per concentration (n=30 eggs/concentration). Egg hatching, mortality, pupation, and eclosion were assessed every 24 hours for 27 days using the methods detailed in [29]. Mean pr(mortality), days to pupation, and days to eclosion were calculated for each concentration and a three-parameter sigmoid curve was fitted to the data to assess LC_50_ prior to eclosion.

### Testing for a change in severity of mannitol’s developmental effects when delaying introduction to larvae by 72 hours

Groups of approximately 100 mixed male and female wild-type flies raised on standard media were placed in each of 10 egg laying chambers. At the end of four hours, eggs were collected and plated on 0M control translucent media where they were raised for 72 hours. After 72 hours, five larvae were plated per treatment petri dish (using translucent media), with the mannitol concentrations from 0 to 0.8 M, at 0.4 M increments. Six petri dishes were used per concentration (n=30 eggs/concentration). Larval mortality, pupation, and eclosion were assessed every 24 hours for another 22 days. Mean percent mortality, days to pupation, and days to eclosion were calculated for each concentration.

### Testing effects of mannitol on oviposition choice

An oviposition choice test [29] was used to assess differences in egg laying choice between control and 0.5M mannitol media (control media, with the addition of mannitol). Groups of ten, 0-24 hour old adult male and ten, 0-24 hour old female flies were reared on standard media for 72 hours before being transferred to choice arenas. The arenas consisted of two vials on their sides, one containing control media and the other containing 0.5 M mannitol media, connected by a plug with a 1.5 cm diameter central tube. Flies were observed moving freely between vials. Six treatment arenas contained a choice between 0.5 M and 0 M mannitol, while six control arenas contained a choice between two 0 M mannitol media.

After 72 hours on standard media, 10 male and 10 female flies were transferred to the paired vial set-ups, with 5 female and 5 male flies placed on either side of the plug. Flies were moved to a new media arena every 24 hours and eggs laid on each media were counted daily for three days (n=7,120 eggs). Vial orientation within the incubator was rotated once per day.

### Statistical analyses

Analyses were performed using SPSS v. 24, Sigmaplot v 12.5, and Graphpad v. 8.0.0 [36–38]. The effects of mannitol introduction to larvae on adult body size were analyzed using Kruskal-Wallis test with Dunn’s multiple corrections for each sex. A 2-way ANOVA was used to look for an interaction effect between sex and mannitol concentration on body size. A linear regression was fitted to the data for each sex across concentrations, and the slopes and intercepts were compared in Graphpad to assess if sexes differed in body size and in the degree of mannitol’s effect on their body size.

Effects of eclosion day on male or female body size within a concentration were assessed using linear regressions in GraphPad, to understand the effects of mannitol in individuals that are more or less delayed in their development within a concentration and sex. This allowed us to look for any effect of day-based sampling bias, as we did not measure every emerging adult’s body size, but only two per day of each sex in each vial. There was no significant trend within each pair of concentration and sex (e.g. 0 M + females) of emergence day on body size, except in 0.4 M males, indicating that flies emerging earlier and later within a concentration were not differently affected by mannitol and reducing the likelihood of day-based sampling bias on our results (S1 Fig; 0 M-female, F=0.47, p=0.50; 0 M-male, F=3.52, p=0.07; 0.4 M-f, F=0.80, p=0.37; 0.4 M-m, F=10.51, p=0.002; 0.8 M-f, F=0.16, p=0.69; 0.8 M-m, F= 2.00, p=0.16). The slopes of the regressions across all six concentration-sex pairs were not significantly different from one another (F=0.53, p=0.75).

Larval mortality data across mannitol concentrations at 48 hours, 72 hours, and pre-eclosion was assessed using survival analyses in SPSS [39], with subjects living to the end of the trial or eclosed included in the analysis as right-censored values on the final day of that test (48 hours, 72 hours, and the last day of the trial respectively). Pupae that had not eclosed after at least six days at the end of the trial were marked as ‘dead’ on the final day of the trial (day 27). Differences in survival distributions across concentrations were tested using pairwise log-rank Mantel Cox tests. Three-parameter, best-fit sigmoidal function LC_50_ curves for larvae at 72 hours, pre-pupation, and pre-eclosion were generated in Sigmaplot. To analyze any effects on survival of delaying the introduction of mannitol to larvae by 72 hours, we used a pairwise log-rank Mantel Cox test (with subjects eclosed before the end of the trial included as right-censored values on day 25, and pupae that had not eclosed marked as ‘dead’ on the final day).

To analyze developmental delays across concentrations, we used a one-way ANOVA with Tukey’s multiple comparisons test in Graphpad. To analyze differences in time from pupation to eclosion, a one-way ANOVA with Tukey’s multiple comparisons was used. To analyze any phenotypic effects on pupation/eclosion time across replicates (n=6/concentration) by delaying the introduction of mannitol to larvae by 72 hours, we used a 2-way ANOVA and Tukey’s multiple comparisons Tests in Graphpad. Differences in the number of larvae that pupated, but did not eclose, across concentrations in the delayed-introduction treatments were analyzed using Fisher’s exact tests in Graphpad.

Oviposition choice was tested using a chi square against a 50-50 population, and against our control-control vial populations.

## Results

### Effects of larval ingestion of mannitol on adult body size

Adult female body size decreased as mannitol concentration increased, with 0.8 M emerging adults having smaller body sizes than 0 M or 0.4 M emerging adults (Fig 1, Dunn’s: 0 M-0.8 M, Z=4.44, p<0.0001; 0.4 M-0.8 M, Z=2.59, p=0.029; 0 M-0.4 M, Z=2.12, p=0.10). Male body size also decreased as mannitol concentration increased, with 0.8 M and 0.4 M emerging adults having smaller body sizes than 0 M emerging adults (Fig 1, Dunn’s: 0 M-0.8 M, Z=4.77, p<0.0001; 0 M-0.4 M, Z=4.12, p=0.0001; 0.4 M-0.8 M, Z=0.88, p>0.99). For females, the linear regression of mannitol concentration on body size was y=-0.04930x+1.022 (F=21.7, p<0.0001, R^2^=0.12); for males, y=-0.04644x+0.8992 (F=26.90, p<0.0001, R^2^=0.14). The slopes did not differ between males and females (F=0.04, p=0.84) indicating increasing mannitol concentration did not affect one sex’s body size differently than the other (2-way ANOVA: interaction effect, F=1.07, df=2, p=0.34). The intercepts were significantly different (F=792.6, p<0.0001) indicating females had larger body sizes than males at all concentrations (2-way ANOVA: sex, F=769.2, df=1, p<0.0001).

**Fig 1.**
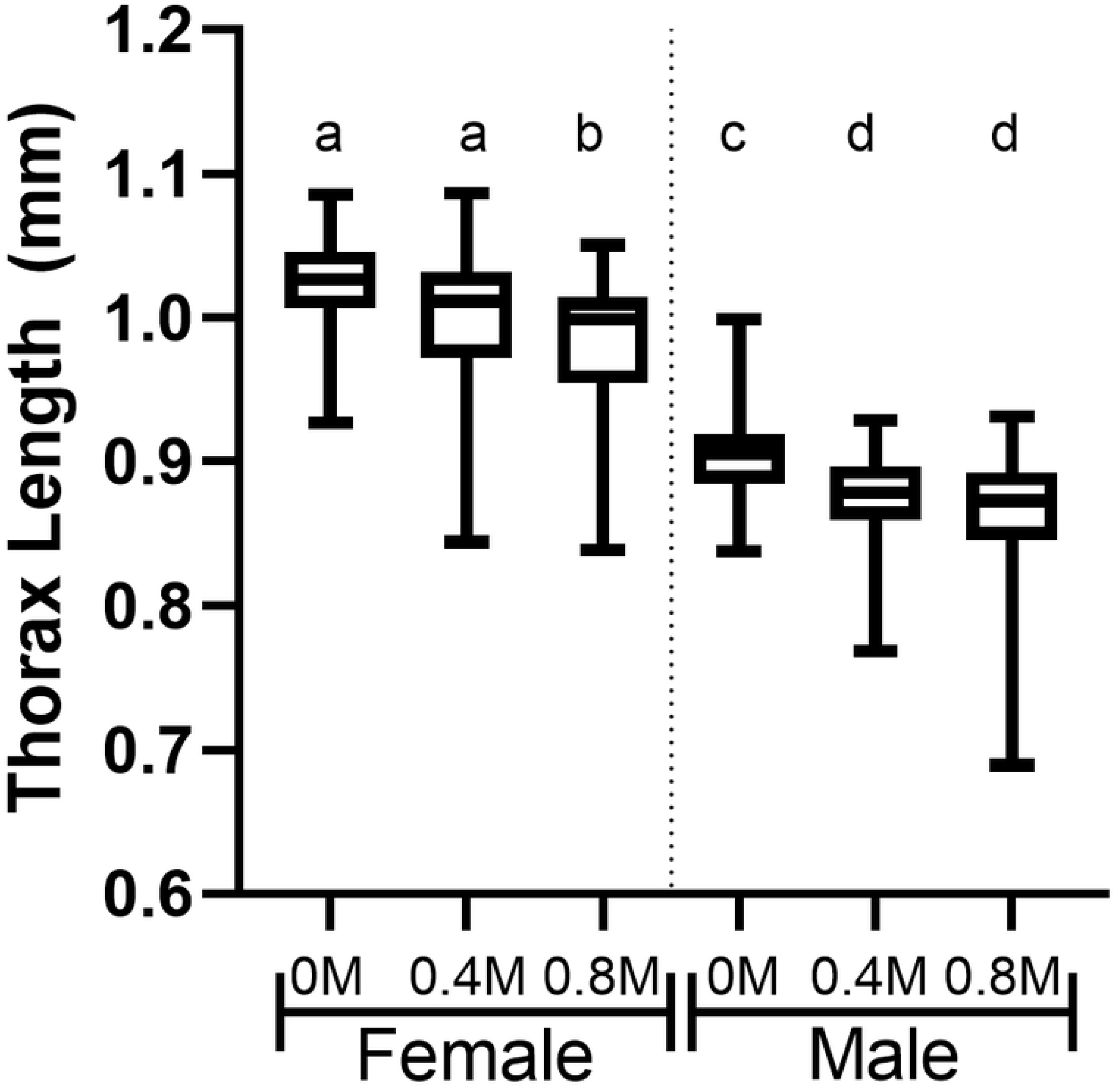
Concentration-dependent decreases in body sizes of adult *D. melanogaster* fed mannitol as larvae. Boxplots showing thorax lengths of males and females across increasing concentrations of mannitol; ingesting increasing mannitol concentration as larvae significantly decreases thorax lengths in emerging adults. Letters indicate significant differences between treatments (Dunn’s: p<0.05). Linear regressions show larval ingestion of increasing mannitol concentrations decreases emerging adult thorax lengths in males and females [females: y=-0.04930x+1.022 (F=21.7, p<0.0001, R^2^=0.12); for males, y=-0.04644x+0.8992 (F=26.90, p<0.0001, R^2^=0.14)].

### Concentration-dependent developmental delay prior to the onset of pupation and reductions in survival

#### Developmental delay

Time to pupation was significantly increased in the 0.4 M, 0.6 M, and 0.8 M conditions as compared to controls (Fig 2, ANOVA with Tukey’s: 0.4 M, q=8.61, p<0.0001; 0.6 M, q=14.35, p<0.0001; 0.8 M, q=8.97, p<0.0001), but not the 0.2 M condition (q=3.15, p=0.18). Time to adult eclosion was significantly increased in all the treatment conditions as compared to controls (ANOVA with Tukey’s: 0.2 M, q=4.11, p=0.04; 0.4 M, q=8.96, p<0.0001; 0.6 M, q=14.85, p<0.0001; 0.8 M, q=11.52, p<0.0001). However, the time between pupation and eclosion was not significantly different from controls in any mannitol treatment (S2 Fig, ANOVA: F=1.04, p=0.39), indicating the major cause of eclosion delay was a delay in the onset of pupation.

**Fig 2.**
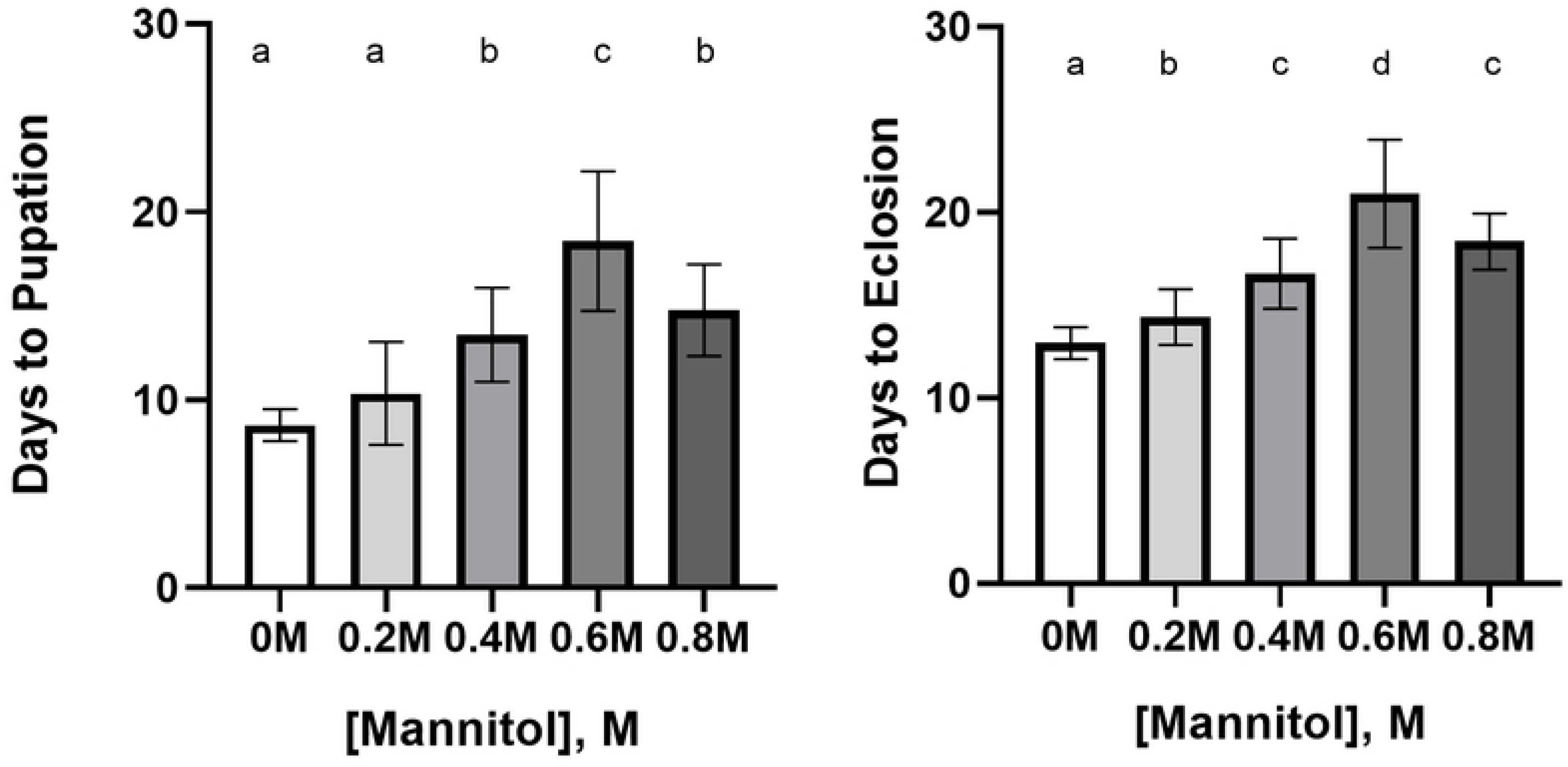
Concentration-dependent developmental delay in *D. melanogaster* larvae fed increasing concentrations of mannitol. **(left)** Time to pupation in *D. melanogaster larvae* was significantly increased in 0.4 M-0.8 M conditions as compared to 0.2 M and control conditions. Letters indicate highly significant differences between concentrations (ANOVA with Tukey’s, p<0.05). **(right)** Time to eclosion in *D. melanogaster* pupae was significantly increased in 0.2 M-0.8 M conditions. Letters indicate significant differences between concentrations (ANOVA with Tukey’s, p<0.05). Error bars represent one standard deviation.

#### Reduced survival

We next assessed the effect of mannitol on *D. melanogaster* larval and pupal mortality. Mortality was concentration dependent for *D. melanogaster* larvae and pupae when assessed prior to eclosion, with 0.4 M, 0.6 M, and 0.8 M treatments showing a significant difference from the control (S3 Fig, Mantel-Cox: 0.2 M, X^2^=0.28, p=0.60; 0.4 M, X^2^=9.40, p=0.002; 0.6 M, X^2^=23.53, p<0.001; 0.8 M, X^2^=19.41, p<0.001).

Highly significant differences in larval mortality occurred as early as 48 hours after egg laying in the 0.6 M and 0.8 M (Fig 3, Mantel-Cox: 0.6 M, X^2^=5.24, p=0.022; 0.8 M, X^2^=10.39, p=0.001) and 72 hours after egg laying in the 0.4 M, 0.6 M, and 0.8 M (Mantel-Cox: 0.4 M, X^2^=4.47, p=0.035; 0.6 M, X^2^=11.81, p=0.001; 0.8 M, X^2^=11.88, p=0.001).

**Fig 3.**
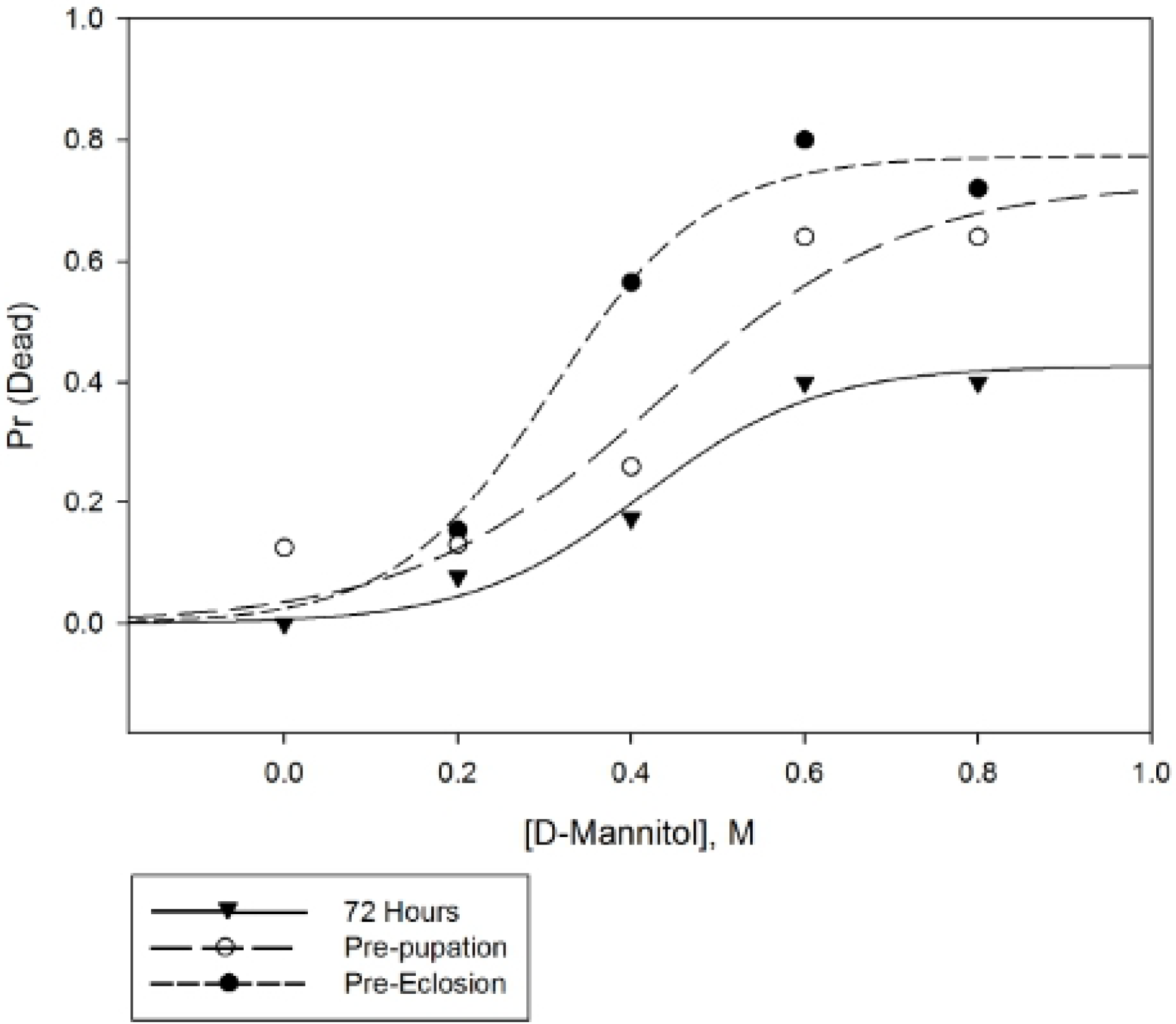
Proportion of larvae dead after mannitol ingestion at different time points during development. Proportion of *D. melanogaster* larvae dead at 72 hours after egg lay, prior to pupation (inclusive of deaths at 72 hours), and prior to eclosion (inclusive of 72 hour and prior to pupation deaths), across increasing concentrations of mannitol. The three-parameter best-fit sigmoidal functions are shown, and the function for pre-eclosion mortality was used to calculate the LC_50_ for *D. melanogaster* prior to eclosion (0.36 M mannitol).

The best-fit sigmoidal curve for pre-eclosion LC_50_ data was:

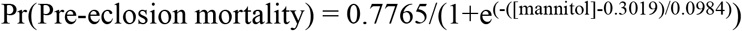

This curve was a significant fit to the data (Fig 3; R^2^=0.96, p=0.039) and using the equation we found the pre-eclosion LC_50_ to be 0.36 M mannitol.

In 0.4 M and 0.6 M treatments, there were significant increases in the proportion of larvae that died prior to eclosion but after pupation, compared to both 0 M (Fig 6, Fisher’s: 0.4 M, p=0.0015; 0.6 M, p=0.0046) and 0.2 M (0.4 M, p=0.0023; 0.6 M, p=0.0062); 0 M and 0.2 M were not different from one another (p>0.99). 0.8 M treatments were not significantly different, but this may be an effect of small sample size due to decreased survival during the larval stage in this treatment (n=9 pupae, Fisher’s: 0 M, p=0.08; 0.2 M, p=0.09).

### Concentration-dependent reduction of mannitol’s developmental effects by delaying mannitol introduction to larvae for 72 hours

#### Partial rescue of developmental delays

Introducing larvae to mannitol after 72 hours [72-hour plates] significantly decreased pupation and eclosion times in the 0.4 M treatment (Fig 4a; ANOVA with Tukey’s, pupation, q=12.71, p<0.0001; eclosion time, q=7.94, p<0.0001), and the 0.8 M treatment (pupation time: q=7.02, p<0.0001; eclosion time: q=5.23, p=0.0047) as compared to plates where larvae were fed the same concentration of mannitol from hour 0 after egg lay.

**Fig 4.**
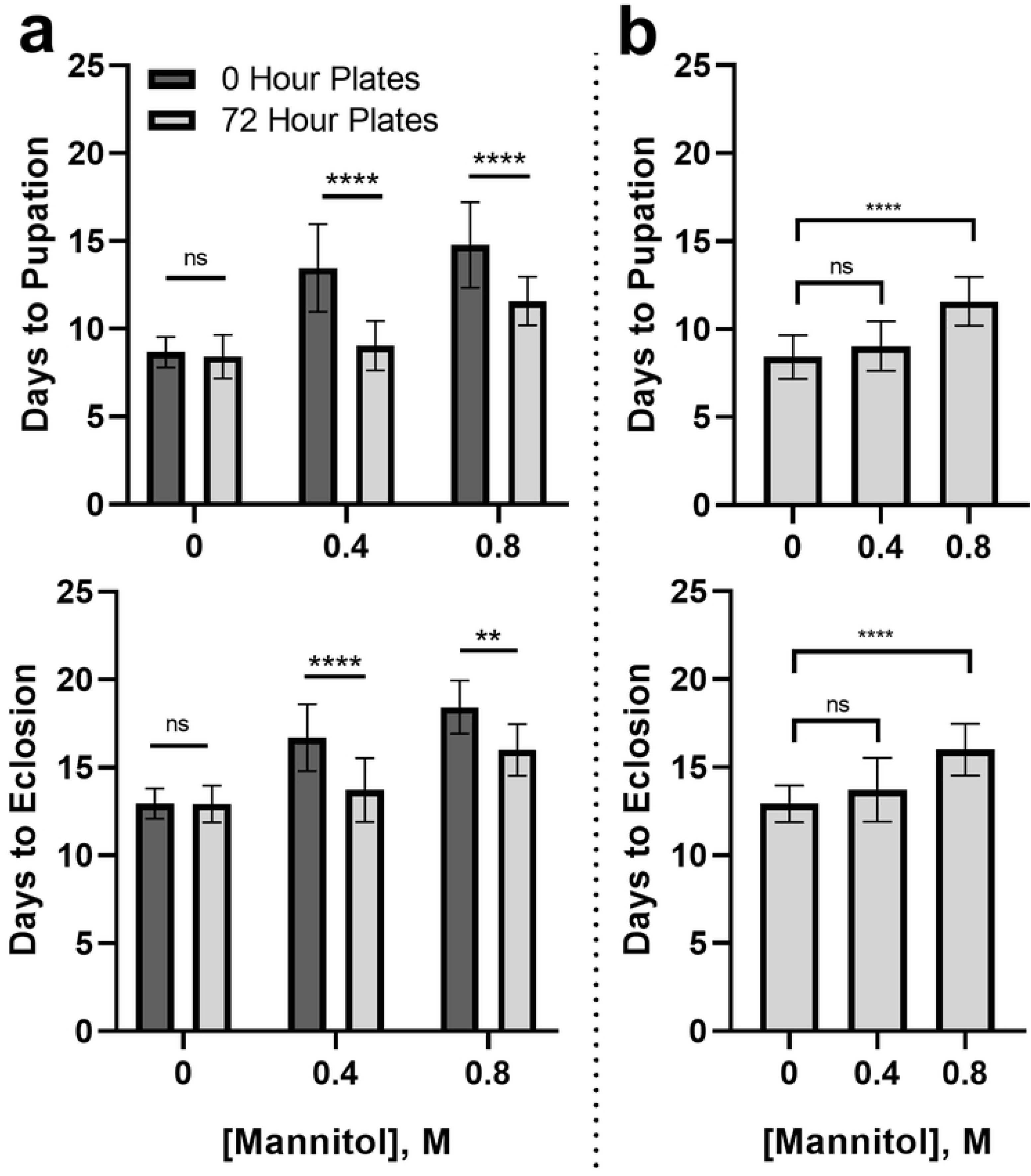
Differences in developmental delay when mannitol introduction is introduced after 72 hours. **(a)** Pupation and eclosion times in *D. melanogaster* larvae were significantly decreased in 0.4 M and 0.8 M conditions when larvae were first placed on mannitol 72 hours after egg lay. Stars indicate significant differences between 0 hour and 72-hour plates (Tukey’s, ns=not significant, **=p<0.01, ***=p<0.001). Error bars represent one standard deviation. **(b)** Pupation and eclosion times were not significantly different between 0 M and 0.4 M treatments when larvae were first placed on mannitol 72 hours after egg lay; larvae fed 0.8 M mannitol after 72 hours still had prolonged pupation and eclosion times. Stars indicate significant differences between 0 hour and 72-hour plates (Tukey’s, ns= not significant, ****=p<0.0001). Error bars represent one standard deviation.

Pupation and eclosion times were no longer significantly different from 0 M conditions in the 0.4 M 72-hour plates (Fig 4b; ANOVA with Tukey’s, pupation, q=2.00, p=0.72; eclosion, q=2.82, p=0.35). Pupation and eclosion times were still significantly longer than controls in 0.8 M 72-hour plates (pupation: q=9.30, p<0.0001; eclosion: q=9.20, p<0.0001).

#### Partial rescue of larval survival

Waiting 72 hours before introducing larvae to mannitol media also significantly increased survival to eclosion across replicates at 0.4 M and 0.8 M (Fig 5a, Mantel-Cox: 0.4 M, X^2^=8.91, p=0.003; 0.8M, X^2^=6.80, p=0.009). In the 0.4 M 72-hour plates, survival was no longer significantly different from 0 M treatment (Fig 5b; X^2^=0.00, p=0.986). The 0.8 M 72-hour plates treatments were still significantly different from 0 M (X^2^=8.03, p=0.005).

**Fig 5.**
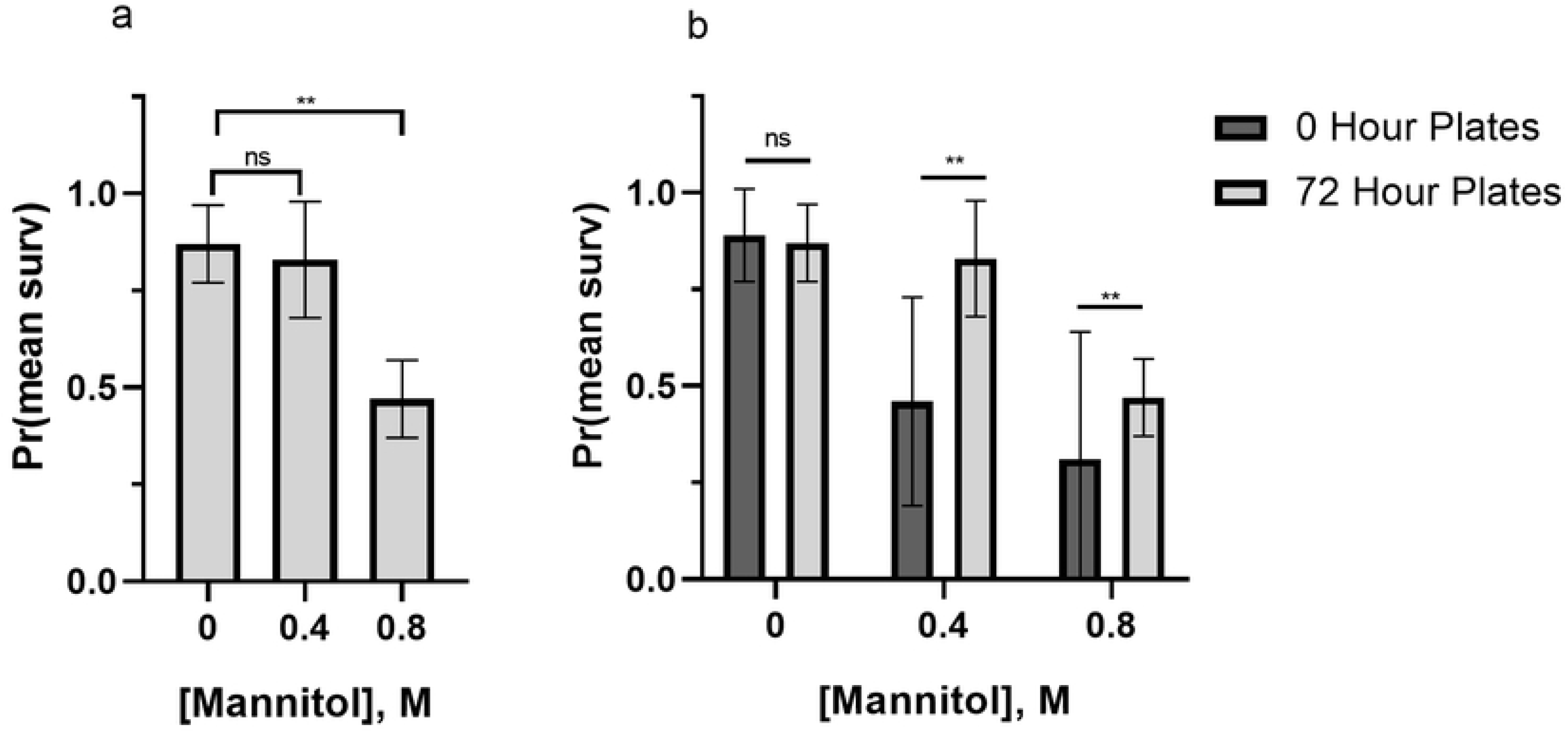
Concentration-dependent partial rescue of survival when mannitol is introduced after 72 hours. **(a)** When mannitol introduction to *D. melanogaster* larvae is delayed by 72 hours, 0.4 M and 0 M treatments no longer differ in their survival while 0.8 M treatments still have significantly decreased survival compared to controls. Stars indicate significant differences between 0 hour and 72-hour plates (Mantel-Cox, ns=not significant, **=p<0.01). Error bars represent one standard deviation. **(b)** Pre-eclosion survival was significantly increased in 0.4 M and 0.8 M conditions when larvae were first placed on mannitol media after 72 hours instead of at hour 0 (egg lay). Stars indicate significant differences between control and 72-hour treatments (Mantel-Cox, ns=not significant, **=p<0.01). Error bars represent one standard deviation. The percent of pupae that did not eclose significantly decreased in 0.4 M treatments when mannitol introduction was delayed by 72 hours, but no significant difference was found between 0 hour and 72 hour mannitol introduction in 0.8 M treatments (Fig 6, Fisher’s: 0.4 M, p=0.017; 0.8 M, p>0.99). The percent of pupae that did not eclose in 0.4 M 72-hour plates was not significantly different from 0 M controls (Fisher’s: p>0.99).

**Fig 6.**
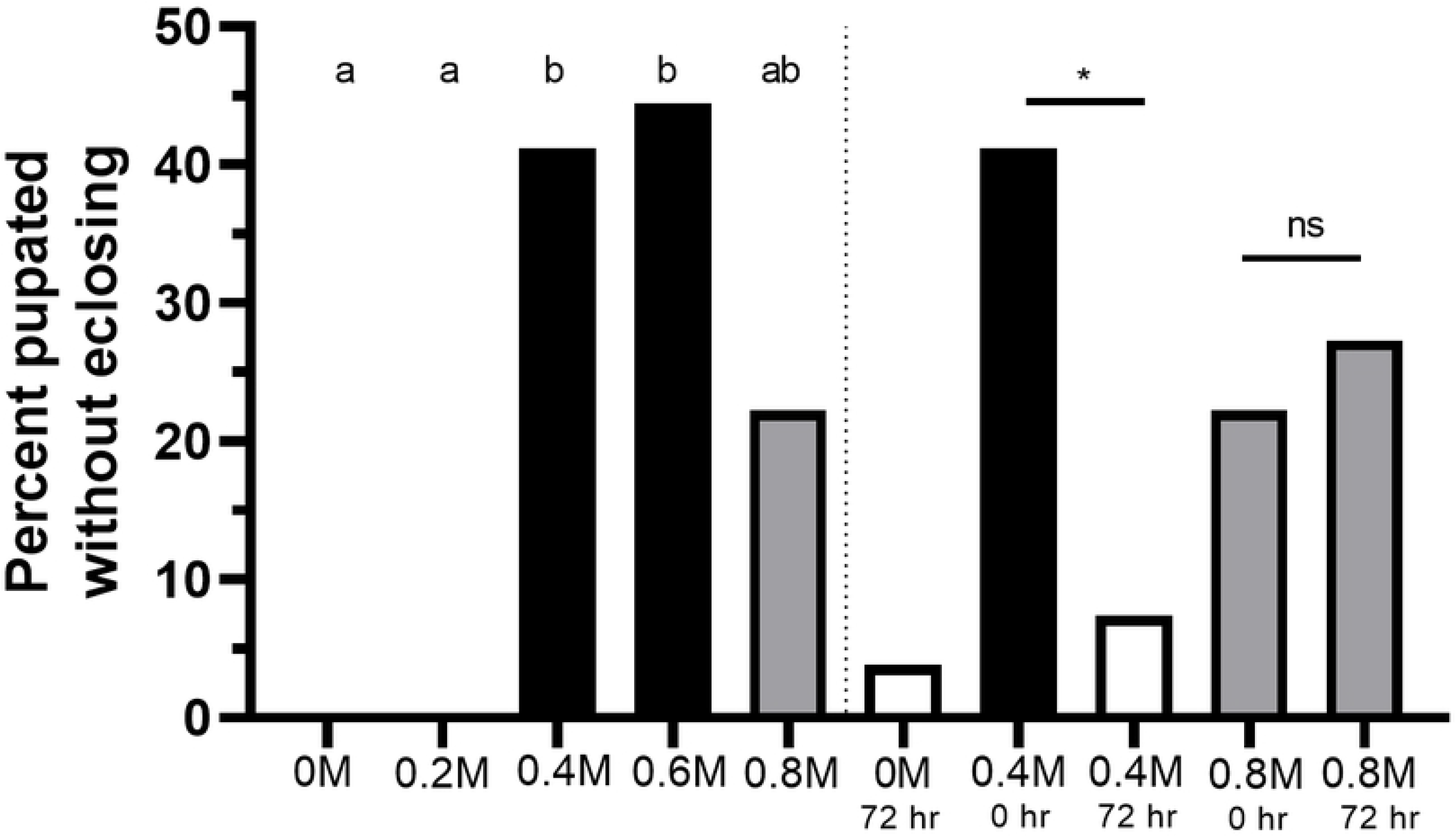
Concentration-dependent eclosion failure, and change in eclosion failure due to delayed mannitol introduction, across increasing concentrations of mannitol. Percent of larvae that pupated but failed to eclose across increasing concentrations of mannitol (0 M-0.8 M) and in 0.4 M and 0.8 M 72-hour plate treatments. Letters indicate highly statistically significant differences between treatments (Fisher’s: p<0.01). Stars indicated significant differences between 72 hour and 0 hour plates of the same concentration (Fisher’s: ns=not significant, *=p<0.05).

### Mannitol avoidance in female oviposition choice assays

The number of eggs laid in the 0.5 M mannitol vials of the mannitol-control choice arenas was significantly lower than expected when compared to a 50-50 population of the same number, or to either side of the control-control arenas (Fig 7, Chi square: 0.5 M vs 50-50, X^2=^514.0, p<0.0001; 0.5 M vs 0 M left side, X^2^=600.8, p<0.0001 or vs 0 M right side, X^2^=392.3, p<0.0001). 24.7 % of the total eggs were laid in the mannitol side.

**Figure 7.**
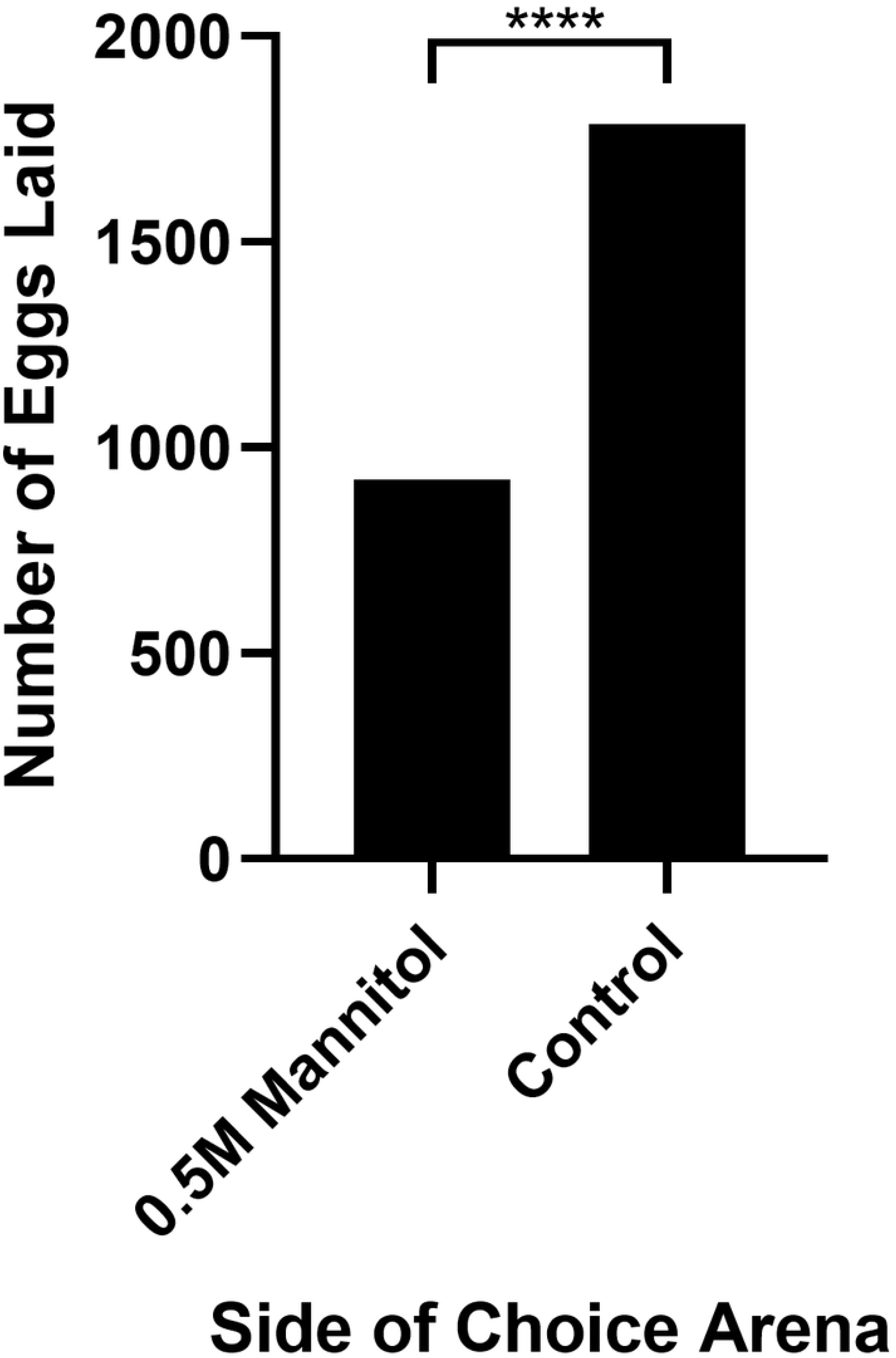
Adult females avoid laying eggs on mannitol foods in choice arenas. Females laid more eggs on control media than 0.5 M mannitol media in mannitol-control choice arenas (n=7,120 eggs; Chi square, ***= p<0.0001).

## Discussion

Several recent studies support the idea that the positive relationship between body size and developmental duration can be reversed due to variable diets, even across different insect taxa [17,40–42]. The phenotypic effects of a mannitol diet on this relationship in *D. melanogaster* are similar to the effects of high-sugar diets, produced via disrupting the insulin-signaling pathway [10,12,17,18]. Mannitol increased *D. melanogaster* developmental duration and decreased emerging adult body size in a concentration-dependent manner. Increased developmental duration was a result of delayed onset of pupation, not prolonged pupal metamorphosis. Stage of larval development when mannitol was introduced (first or third larval instar) and mannitol concentration influenced the severity of mannitol’s phenotypic effects; 0.4 M mannitol introduced at 72 hours no longer affected development time or survival, but 0.8 M mannitol still had significant, if lessened, effects. These phenotypic effects are consistent with those of high-sugar diets that generate smaller adult body sizes and prolonged development prior to the onset of pupation [10,17,18]; as with mannitol, the high-concentration sugar diets have stronger effects earlier in larval development (prior to the third instar) [10]. Females avoided mannitol foods as oviposition substrates despite the heightened carbohydrate concentration, indicating that mannitol (a product of microbial fermentation) may provide important information about oviposition site quality to females [16].

Models of the independent effects of growth rate, critical weight, and the interval to the cessation of growth on the relationship between body size and developmental duration in *Manduca sexta* indicate that variation in growth rate can lead to a negative relationship (decreasing body size while increasing developmental duration), while variations in interval to the cessation of growth and critical weight generally lead to positive relationships [4,43]. Critical weight typically occurs directly after the second molt in *D. melanogaster*, at approximately 72 hours post-hatching [2,31]. Mannitol, when given at high concentrations only after 72 hours of development, still increased *D. melanogaster* developmental times, making it unlikely that mannitol decouples the positive relationship between body size and developmental duration via altering critical weight. Instead, mannitol is likely impacting growth rate and/or the interval to cessation of growth, potentially by disrupting the insulin/TOR signaling pathway which is responsible for regulating these variables in *D. melanogaster* [44].

In *D. melanogaster*, a carbohydrate-rich diet led to delays in eclosion and smaller pupal case sizes [17]. Extremely high sugar (e.g., 1 M sucrose) diets produced insulin resistance, leading to smaller wandering third instar larvae and smaller eclosed adults irrespective of protein availability, the sugar used, or osmolarity of the food medium during development [10]. In addition, high-sugar feeding led to dramatic delays in pupation [10,12], similar to what we saw in our 0.4 M - 0.8 M mannitol treatments. Delays in eclosion due to high-sugar diets affecting the insulin-signaling pathway cause delayed onset of pupation, not prolonged metamorphosis [18]; again, this is the same phenotype we saw when larvae were fed mannitol diets.

Feeding third instar larvae high-sugar diets for just 12 hours produced similar transcriptional effects in genes associated with glucose transport and metabolism, lipid synthesis and storage, trehalose synthesis and stability, and oxidative stress compared to larvae being fed high-sugar diets since egg lay, just with lower fold-changes in expression [10]. We observed that third instar larvae fed 0.4 M mannitol had normal developmental durations, while third instar larvae fed 0.8 M mannitol had only partially alleviated mannitol’s developmental effects. Lower, but still significant, levels of gene expression changes in third instar larvae due to the introduction of high-sugar diets may explain how mannitol’s developmental effects were not completely restored to normal at the highest concentrations (0.8 M), even when mannitol introduction was delayed to the third instar.

Because concentrations of all non-mannitol carbohydrates were kept the same in larval foods, *D. melanogaster* would need to be able to metabolize mannitol in order for it to increase levels of trehalose in the hemolymph like the other, metabolizable sugars (glucose and sucrose) used in previous studies. No studies have examined if *D. melanogaster*, or its common gut microbes, can metabolize mannitol, but recent work on another insect, *Tribolium castaneum*, shows that adult females have higher trehalose levels in the hemolymph after feeding on mannitol [26]. Circulating trehalose is responsible for TOR activation in *D. melanogaster* fat bodies, contributing to cell growth during development; mannitol’s catalysis to trehalose may be responsible for mediating its effects on growth rate and the interval to the cessation of growth via the insulin/TOR signaling pathway similar to other carbohydrates [11,45].

The insulin/TOR signaling pathway regulates development in response to nutrients, and disruptions of this pathway are known to affect body size and developmental duration [19,44,46,47]. Genetic defects in the insulin signaling pathway (including dILPs, *Drosophila* insulin-like proteins), reduced insulin receptor activity, the inhibition of DREF (DNA replication-related element-binding factor), or disruptions in TOR activation, can cause long development times and smaller-bodied flies, similar to the adverse mannitol developmental effects we observed in larvae [9,19,48–53]. High-sugar diet fed larvae experience reduced growth due to insulin signaling resistance, even when dILP levels are increased, while reductions in nutrition can simply prevent the release of dILPs, thereby reducing growth [10,44]. dILP expression in response to a mannitol diet should be explored to better understand if disruptions to the insulin/TOR signaling pathway mediate mannitol effects on larval body size, growth, and developmental duration, especially given the similarity between the adverse effects of mannitol and the adverse effects of high-carbohydrate diets on *D. melanogaster* larvae.

Proper growth during development can also influence survival to, and in, adulthood [28]. High sugar diets cause mortality in larvae of numerous fly species, including *D. melanogaster* and *Drosophila mojavensis* [12,18]; We found that mannitol causes mortality in *D. melanogaster* larvae after 48 hours in a concentration-dependent manner, with an LC_50_ of 0.36 M. In addition, of the larvae that pupated in the 0.4 M and 0.6 M treatments, a significant number of them failed to eclose. Adverse mannitol effects were ameliorated in third instar larvae fed 0.4 M mannitol only after 72 hours of development, but not in larvae fed 0.8 M mannitol after 72 hours, indicating that mannitol’s effects depend on both the developmental stage and concentration at which it is introduced.

Alternatively, starvation and/or osmotic stress could be potential mechanisms for mannitol’s effects on larval survival. However, mannitol’s effects are unlikely to be related strictly to starvation given mismatches between starvation phenotypes and our results. Post-critical weight starvation causes accelerated emergence (our 72 hour plates saw normal or delayed emergence) while pre-critical weight starvation causes developmental delay but normal adult body sizes (unlike our smaller adults) [54]. Simply reducing nutritional availability throughout development generates smaller adult body sizes, but no change in survival through eclosion [55]. For these reasons, as well as the fact that all mannitol-fed larvae received the same basic nutrients as controls, we consider reduced nutritional availability to be an unlikely driver of mannitol’s lethality.

Mannitol may also be acting as an osmotic stressor to larvae, as mannitol is known for its diuretic effects [56–58]. Other species (including the Dipteran, *Aedes aegypti*) exhibit longer development times, decreased body size, and/or reduced survival in osmotically stressful environments [59–62]. *D. melanogaster* larvae have a severe aversive reaction to high concentrations of osmotically stressful substances like salt [63]. This aversive reaction is coupled with larvae decreasing their food intake [63], and decreases of 30 % in larval mass at the third instar [64]. Extremely high sugar diets (20% sucrose) have been shown to decrease feeding in *D. melanogaster* larvae [12] and decreased feeding at the highest mannitol concentrations may explain the atypical trends of survival and developmental duration in our 0.8 M conditions (where 0.6 M mannitol often had slightly more adverse effects than 0.8 M mannitol). However, it should be noted that *D. melanogaster* larvae have excellent osmoregulatory ability and other *Drosophila* species’ larvae have been found living in abundance in osmotically stressful, high sugar environments [65,66].

Our oviposition choice data shows that females avoid laying on mannitol media, even when the media has more abundant carbohydrates, suggesting they may be able to perceive the presence of mannitol at concentrations of 0.5 M and above. As a product of microbial fermentation in many microorganisms [21,23], mannitol may indicate important information about the quality of oviposition locations to females, especially given that females pick nutritional compositions of pre-rotting fruit over a composition more similar to currently-rotting fruit to lay their eggs (despite these environments being suboptimal for larval performance at the time of egg lay; [16,67]). Adult body size affects individual fecundity [68,69], so female *D. melanogaster* should attempt to avoid oviposition sites that reduce the body size and survival of offspring.

This study is the first to examine the effects of mannitol on development in any species of holometabolous insect. In the sweet potato whitefly (*Bemisia tabaci*), mannitol was found not to be lethal to nymphs, only adults, at a concentration of 10% [70]. Given mannitol’s vastly different effects on adults of different species (from nutritive to lethal), more work should be done to understand mannitol’s effects on development across taxa [20,26,27,70–74]. This may further our understanding of how species differ mechanistically in their responses to this polyol, particularly important since adverse mannitol developmental effects closely align with the phenotypic effects of high-sugar diets on *D. melanogaster* larvae, mediated via the insulin-signaling pathway.

A single, genetically variable insulin signaling pathway regulates growth, reproduction, longevity, and metabolism in all insects, and contains conserved elements across all animals [75,76]. This pathway is involved in numerous examples of environmentally-driven polyphenisms generated during insect development, including caste differentiation in social insects, as well as geographically- and nutritionally-driven morphological variation [77–81]. Genetic and epigenetic differences in this pathway allow species and populations to take unique advantage of environmental variables (e.g. diet, temperature, seasonality), generating phenotypic plasticity in developmental variables and allometric relationships that maximize fitness [79–81]. Mannitol’s developmental effects, if acting through this pathway, provide an opportunity to compare phenotypic variation in response to the nutritional environment, generated by the evolution of insulin signaling genes across species. This study joins a growing body of work indicating that the frequently-cited positive relationship between duration of development and body size within a population can be complicated by environmental variation, particularly via dietary influences on insulin signaling. Our work also suggests that the importance of this variation, and its influence on specific developmental parameters, may change as development progresses past various internal regulatory cues. Mannitol’s effects on development provide a novel paradigm for exploring the environmentally-cued regulation of developmental-physiological relationships in insects.

## Conclusion

Mannitol causes concentration-dependent developmental delays, smaller adult body sizes, and decreased survival in *D. melanogaster* larvae and pupae. These adverse developmental effects can be alleviated at a concentration of 0.4 M if mannitol introduction is delayed until larvae are 72 hours old (approximately L3), but not at a concentration of 0.8 M. Adult females perceive and avoid mannitol (at a concentration of 0.5 M) when given a choice in oviposition site, likely due to mannitol’s adverse effects on offspring survival and in spite of the mannitol food having more abundant carbohydrates. Mannitol at high concentrations may be acting via the insulin signaling/TOR pathway to decouple the typical, direct relationship observed between body size and developmental duration, creating similar effects to other high-carbohydrate diets.

## Acknowledgements

We thank Meghan Campbell, Natalie Carroll, Virginia Caponera, and Lauren Hultgren for assistance caring for and counting larvae. Kaitlin Baudier provided assistance with basic methodological and statistical procedures. Jennifer Viveiros provided support and training in fly husbandry, sexing, and making foods. This work was supported by funds from Drexel University’s Office of Research (to DRM). Work in the Marenda lab is supported by the National Science Foundation, IOS-1856439 (to DRM).

## Supplementary Information

**S1 Fig.** Linear regressions showing effect of emergence day on thorax length in males and females at each concentration. Only 0.4M-males saw a significant decreasing in thorax length as eclosion was more delayed (y=-0.0029x+0.9777; F=10.51, p=0.002, R^2^=0.1534); slopes were not significantly different from one another (F=0.5298, p=0.7537). Error bars represent one standard deviation.

**S2 Fig.** Time from pupation to eclosion did not differ between control and any mannitol treatments (ANOVA, F=1.04, p=0.39). Error bars represent one standard deviation.

**S3 Fig.** Survival plots showing percent survival to eclosion versus post-hatching fly age given control food or foods with increasing concentrations of mannitol (0.2M to 0.8M). Observations were terminated at 27 days after egg laying (n=30eggs/treatment). Highly significant differences (p<0.01) from the control are in black, non-significant differences are in grey.

